# A megatransposon drives the adaptation of *Thermoanaerobacter kivui* to carbon monoxide

**DOI:** 10.1101/2024.09.20.614114

**Authors:** Rémi Hocq, Josef Horvath, Maja Stumptner, Gerhard G. Thallinger, Stefan Pflügl

**Author notes:** Address correspondence to Stefan Pflügl.

## Abstract

Acetogens are promising industrial biocatalysts for upgrading syngas, a gas mixture containing CO, H_2_ and CO_2_ into fuels and chemicals. However, CO severely inhibits growth of many acetogens, often requiring extensive adaptation to enable efficient CO conversion (“carboxydotrophy”). Here, we adapted the thermophilic acetogen *Thermoanaerobacter kivui* to use CO as sole carbon and energy source. Isolate CO-1 exhibited extremely rapid growth on CO and syngas (co-utilizing CO, H_2_ and CO_2_) in batch and continuous cultures (µ_max_ ∼ 0.25 h^−1^). The carboxydotrophic phenotype was attributed to the mobilization of a CO-inducible megatransposon originating from the locus responsible for autotrophy in *T. kivui*. Transcriptomics illuminated the crucial role maintaining redox balance likely plays during carboxydotrophic growth. These novel insights were exploited to rationally engineer *T. kivui* to grow on CO. Collectively, our work elucidates a primary mechanism responsible for the acquisition of carboxydotrophy in homoacetogens and showcases how transposons can orchestrate evolution.

## Introduction

Synthesis gas (syngas), a mixture composed of hydrogen (H_2_), carbon dioxide (CO_2_) and monoxide (CO) in variable proportions^1^, serves as a fuel and chemical precursor for either catalytic or gas fermentation processes. Biomass as a renewably available resource is deemed essential for establishing a global circular carbon economy^2^, and is a promising feedstock to generate sustainable syngas. Industrial syngas fermentation can be efficiently carried out by acetogens^3^, a phylogenetically diverse group of autotrophic bacteria. These microbes naturally ferment H_2_ and CO_2_ through the reductive acetyl-CoA pathway (or Wood-Ljungdahl pathway, WLP)^4^, an ancient pathway that is thought to be intricately linked with the origin of life^5^. Within the WLP, CO plays a critical role as a key intermediate for acetyl-CoA formation. Being a possibly abundant energy source in primordial environments, CO may have been instrumental as an inorganic substrate for early life^6^.

As a prerequisite for syngas fermentation, acetogens often need to undergo adaptation or evolution to tolerate and efficiently utilize CO (a lifestyle also known as “carboxydotrophy”), in addition to H_2_ and CO_2_^7–9^. In acetogens, CO toxicity is thought to be primarily caused by inhibition of hydrogenases crucial for redox metabolism^10^, reducing or even abolishing growth. Among these hydrogenases, the [FeFe] hydrogen-dependent CO_2_ reductase (HDCR) from *Acetobacterium woodii* and to a much lower extent the [NiFe] energy-converting hydrogenase (Ech) from *Thermoanaerobacter kivui* were shown to be inhibited by CO^11,12^. Circumventing this inhibition was demonstrated to be essential to promote growth of *A. woodii* on CO^13^. On the other hand, the CO_2_/CO reduction potential is particularly low (E°’ ∼ –520 mV ^14^), and the resulting low potential electrons stemming from CO oxidation can boost acetogenic metabolism by significantly increasing ATP yields compared to “classical” growth on H_2_/CO_2_^13,15^. In turn, higher ATP yields critically provide the opportunity to generate more ATP-intensive products such as short or medium-chain alcohols from syngas, or directly from CO (from industrial off-gas streams or obtained via CO_2_ electrolysis)^16^.

In the thermophilic acetogen *T. kivui*, previous work suggested that the buildup of CO tolerance and utilization might be linked with evolution occurring at the HDCR level, although mutations arising in CO-adapted clones were not functionally characterized^17^. In addition, adaptation to CO yielded strains with low growth rates on CO as sole carbon and energy source (T_d_ ∼ 40 h / µ_max_ = 0.017 h^−1^)^7^, severely limiting the potential of *T. kivui* for syngas bioprocessing. This slow growth on CO likely stems from the fact that adaptation was performed in medium supplemented with yeast extract, which is actively used as a substrate for growth by *T. kivui*^18^. As a result, selection bias was shown to occur upon adaptation to CO + yeast extract, as evolved mutants strengthened their ability to grow on yeast extract^18^, thereby drastically complexifying the functional analysis of the mechanisms underlying adaptation to carboxydotrophy.

Here, we sought to create a robust chassis for thermophilic bioprocessing of syngas^19^ and to elucidate the cognate mechanisms allowing for CO tolerance and utilization. We combined adaptive laboratory evolution (ALE)^20^, bioprocess engineering and multi-omics data analysis to generate a strain rapidly growing on CO as sole carbon and energy source (T_d_ ∼ 2.8 h / µ_max_ = 0.25 h^−1^), as well as to quantitatively and qualitatively evaluate its physiological, genetic and metabolic features. We show that in this strain, the acquisition of carboxydotrophy is intrinsically linked to the mobilization of a circular, CO-inducible, 86 kb megatransposon originating from the WLP locus. We further speculate that the synergistic up– and down-regulation of central redox metabolic genes is the primary effector of the acquisition of carboxydotrophy, by alleviating CO-mediated ferredoxin overreduction. Finally, we validate this theory by enabling carboxydotrophic growth in the WT by overexpressing the energy-converting hydrogenase Ech2 complex.

## Results

### Wild-type *T. kivui* rapidly adapts to efficiently grow on CO as sole carbon and energy source

*Thermoanaerobacter kivui* DSM 2030 was adapted to carboxydotrophic growth by an ALE approach featuring three phases (Figure 1a). First, the DSM 2030 strain was cultivated on H_2_ and CO_2_ to establish robust autotrophic growth in chemically defined mineral medium (without yeast extract and vitamins), yielding a cell population referred to as WT_pop_. Next, two syngas mixtures containing different concentrations of H_2_, CO_2_, CO and N_2_ (either 58:9:30:3 or 24:21:52:3, 2 bar) were separately used to adapt *T. kivui* to the presence of CO. Growth on CO as sole carbon and energy source (30 % CO, 70 % N_2_, 2 bar) was achieved by using cells adapted to the 52 % CO syngas mixture (52S_pop_). Cells adapted to 30 % CO syngas (30S_pop_) were unable to grow on 30 % CO. CO concentration was gradually increased in 10 % increments to 60 % CO, and finally to 100 % CO (CO_pop_). From the first inoculation on 52 % CO syngas, the CO_pop_ culture (robustly growing on 2 bars 100 % CO) was obtained in 31 generations.

**Figure 1:**
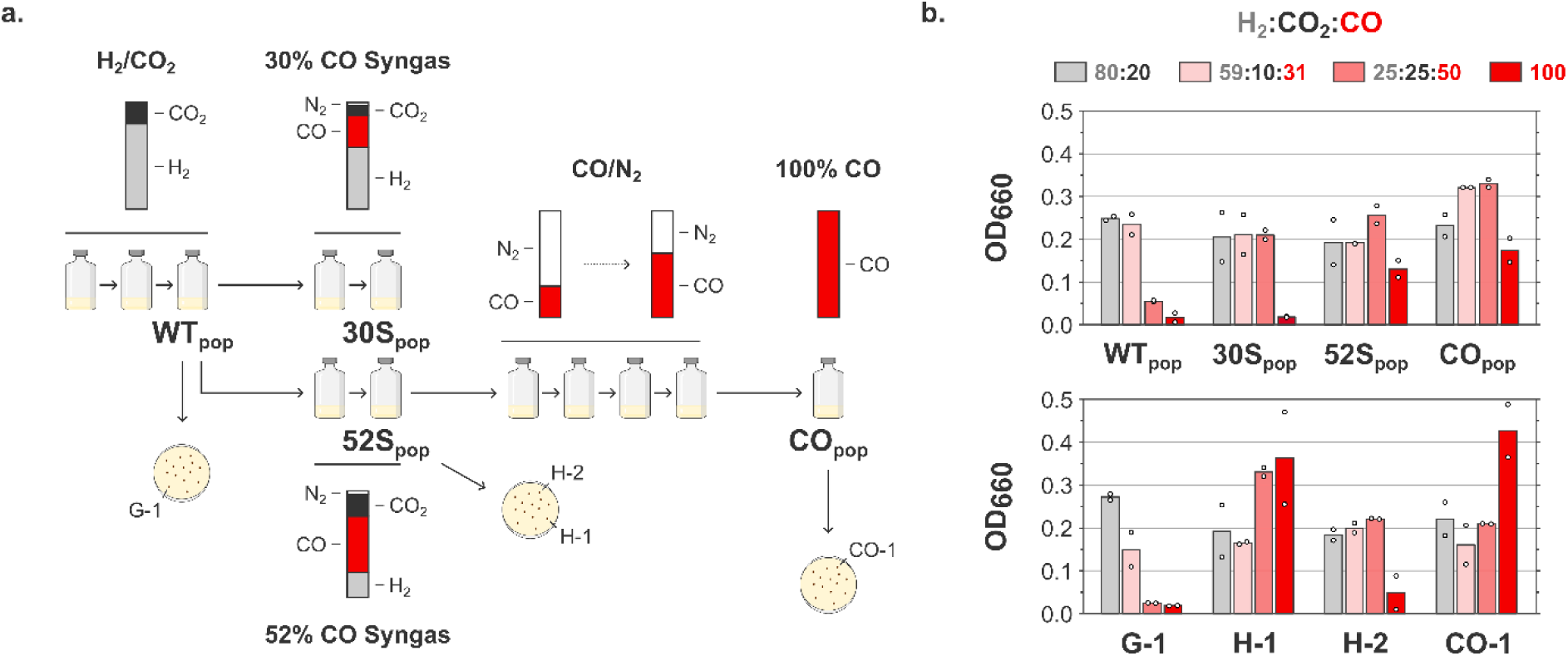
CO adaptation strategy. a) *T. kivui* DSM 2030 was adapted to CO using serial serum bottle cultures with chemically defined mineral medium (without yeast extract and vitamins) under increasing CO concentrations. Clonal strains were isolated on plate from different cultures. b) Biomass formation of adapted populations and clones after 3 days in serum bottles (chemically defined mineral medium without yeast extract and vitamins) under various gas mixtures.

Clonal strains from WT_pop_, 52S_pop_ and CO_pop_ cultures were isolated on plates, and growth of cell populations and clonal strains on various gas mixtures was assessed (Figure 1b). Growth on H_2_/CO_2_ and low CO syngas was possible for all populations and clones. Growth on high CO syngas was observed for all cultures, except for the WT_pop_ and the derived G-1 isolate. 52S_pop_ and related H-1 isolate, as well as CO_pop_ and derived CO-1 clone could all grow on 100 % CO. Strain H-2 derived from the 52S_pop_ population, could interestingly not grow on CO. Collectively, our results indicate that the “pure” carboxydotrophic trait, allowing for growth on CO as sole carbon and energy source, was present already at the 52S_pop_ stage, and therefore likely arose between that culture and the WT_pop_ culture. Only two subculturing steps chronologically separate these cultures (∼ 8-9 generations), suggesting that such a rapid acquisition of the carboxydotrophic trait more likely stemmed from adaptation rather than evolution.

### CO-1 shows high fermentative performance on syngas and CO in bioreactor batch cultivation

To assess the potential of CO-1 for syngas conversion and consolidate its physiological characterization, we evaluated CO-1 fermentative capabilities on two different gas mixtures (CO/H_2_/CO_2_/ N_2_: 52:24:21:3 and 100 % CO) in mineral medium in bioreactor batch cultivations with continuous gassing (Figure 2). The maximum growth rate µ_max_ was particularly high and reached 0.20 ± 0.03 h^−1^ (T_d_ ∼ 3.5 h) and 0.23 ± 0.06 h^−1^ (T_d_ ∼ 3.0 h) on syngas and CO, respectively. High biomass was formed in both conditions (OD_600_ of 2.89 ± 0.72 and 4.85 ± 0.01 for syngas and 100 % CO conditions, respectively) and acetate titers reached high levels (19.9 ± 2.9 and 32.5 ± 7.1 g L^−1^). Importantly, CO, H_2_ and CO_2_ were co-consumed in the syngas condition, which is a critical aspect for syngas bioprocessing. Under 100 % CO, later stages of fermentation show a water-gas shift reaction, resulting in the net production of H_2_ in addition to CO_2_. Together, the results of our batch fermentations confirm that CO-1 is particularly well-suited for syngas and CO conversion.

**Figure 2:**
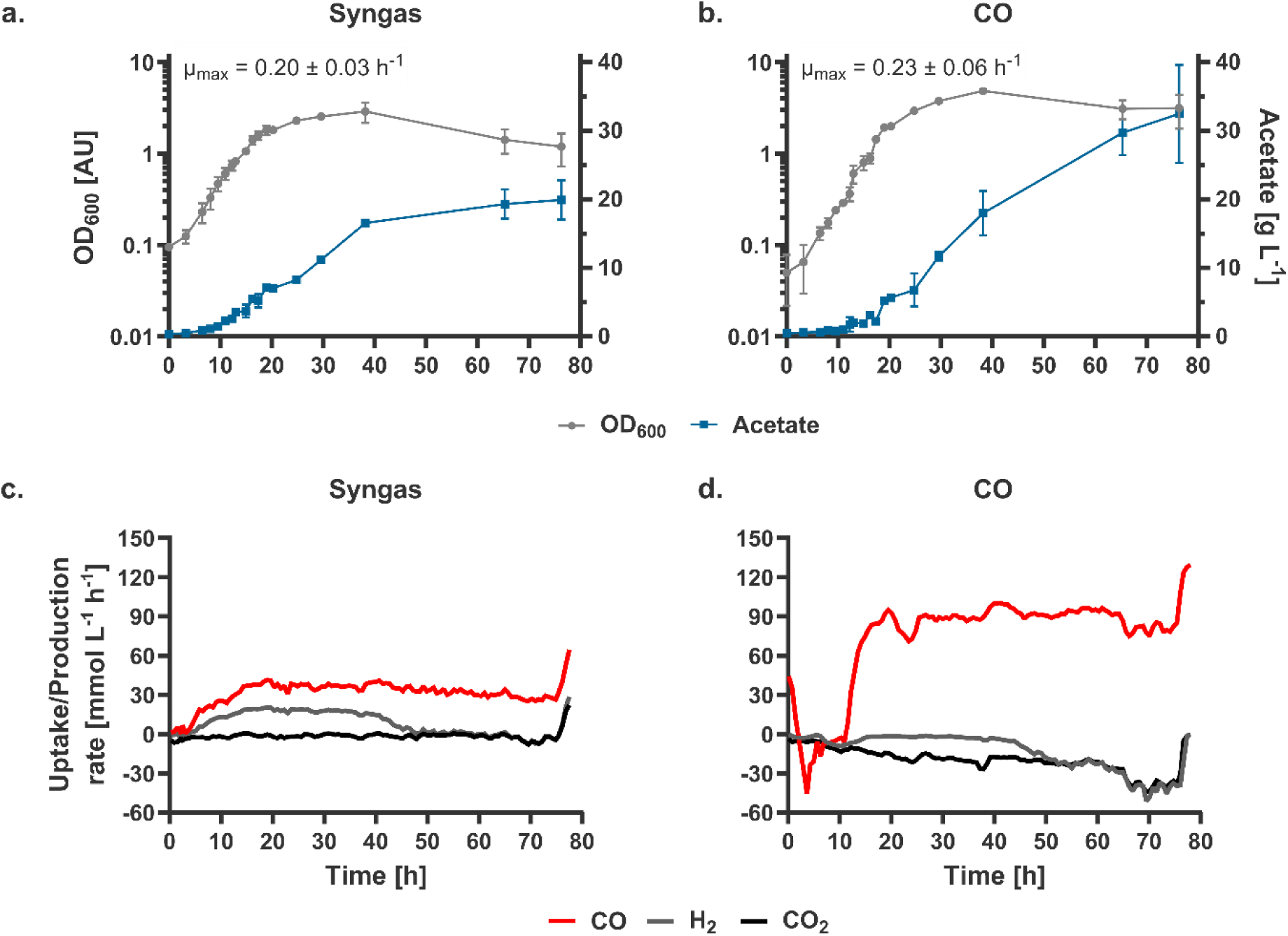
Batch gas fermentation of CO-1 in bioreactor. A synthetic syngas mixture (CO/CO_2_/H_2_/N_2_: 52/21/24/3) and 100 % CO were used as sole carbon and energy source in chemically defined mineral medium (without yeast extract and vitamins). a., b. Biomass (OD_600_), acetate formation and maximum specific growth rate on syngas (a) and CO (b). c., d. Gas uptake (positive) and production (negative) rates on syngas (c) and CO (d). Change of gas uptake/production rates at the end of the experiment due to flushing with N_2_. Error bars represent the standard deviation of duplicate experiments.

### Short-read sequencing analysis reveals only a few SNVs/InDels in the CO-adapted strains

Although *T. kivui* was suspected to have gained the carboxydotrophic trait through adaptation rather than evolution, we submitted the CO-adapted clonal strains for whole genome sequencing (Illumina). These encompass CO-1, as well as two other carboxydotrophic isolates from CO_pop_ (CO-2, CO-3) with a similar phenotype (data not shown). DNA from the non-carboxydotrophic WT_pop_ was sequenced as a reference. As expected, only a few SNVs and InDels were found in the CO-adapted strains (total number across the three strains: 22, Supplementary Table S1). Common mutations in all three strains (Table 1) do not directly affect genes involved in CO metabolism, rendering their potential contribution to the carboxydotrophic phenotype unclear.

**Table 1:** Common mutations found in CO-1, CO-2 and CO-3 strains.

| Position | Type | Locus tag | Effect | Annotation |
| --- | --- | --- | --- | --- |
| 625 773 | Substitution | TKV_RS07415 | D215N (GAT → AAT) | phosphate signaling complex protein PhoU |
| 1 027 204 | Substitution | Intergenic | Intergenic (T → A) |  |
| 1 266 481 | Substitution | TKV_RS04105 | A93P (GCT → CCT) | 4-hydroxy-tetrahydrodipicolinate synthase DapA |
| 1 963 781 | Substitution | TKV_RS10045 | D187E (GAT → GAA) | elongation factor Tu |
| 1 968 524 | Substitution | TKV_RS10070 | A944D (GCT → GAT) | DNA-directed RNA polymerase subunit beta' |

### Short– and long-read sequencing analysis reveals the presence of a large extrachromosomal transposable element in the CO-1 strain

Short-read sequencing can also be used to gain insights into potential large-scale rearrangements, and we therefore performed coverage analysis in the CO-adapted strains. A 2,438 bp deletion was found at positions 233,431 to 235,869, affecting two genes with unknown function as well as another one annotated as N-acetyltransferase (TKV_RS01120 to TKV_RS01130). In addition, two large duplications were found at positions 1,839,957 to 1,900,058 (duplication 1) and 1,908,405 to 1,934,113 (duplication 2), spanning 60,101 and 25,708 bp respectively (Figure 3a). These duplications both display a ∼ 2.2 times higher coverage than the rest of the genome (1765x and 1739x, respectively, compared to 802x). The duplications are separated by 8,348 bp and bear genes crucial for acetogenesis, namely, genes coding for the HDCR, Ech2 and HydABC complexes, as well as a significant part of the WLP locus (Figure 3a). Duplication 1 starts with a gene encoding for an IS1182-type transposase, while duplication 2 ends just before a region rich in tRNA and rRNA genes, which are typical recognition sites for bacterial transposases^21–26^. This disposition as well as the increased coverage strongly suggested that the large genomic region encompassing the WLP locus could be mobilizable by transposition elements. We therefore submitted DNA extracted from CO-1 to long-read sequencing (Oxford Nanopore Sequencing, ONT) to pinpoint the localization of the additional copies of duplications 1 and 2. *De novo* assembly yielded a circular 2,396,992 bp sequence identified as *T. kivui* chromosome (WT chromosome: 2,397,805 bp), as well as an additional circular sequence of 85,811 bp. This extrachromosomal element, designated as Tn_CO-1_, links both duplications as depicted in Figure 3b. Junction sequences linking both duplications are specific to Tn_CO-1_ and primer pairs amplifying these short Tn_CO-1_-specific sequences (which cannot be found on the genome) could therefore be designed (Figures 3a and 3b). Successful PCR amplification was observed for CO-1 gDNA, but not G-1 gDNA, confirming the presence of Tn_CO-1_ *in vivo* (Figure 3c). Additional transposon activity was found when aligning CO-1 and WT chromosomes, with an insertion of an ISL3 family transposase and a translocation of an ISLre2 family transposase, neither affecting genes involved in carboxydotrophy (Supplementary Table S2).

**Figure 3:**
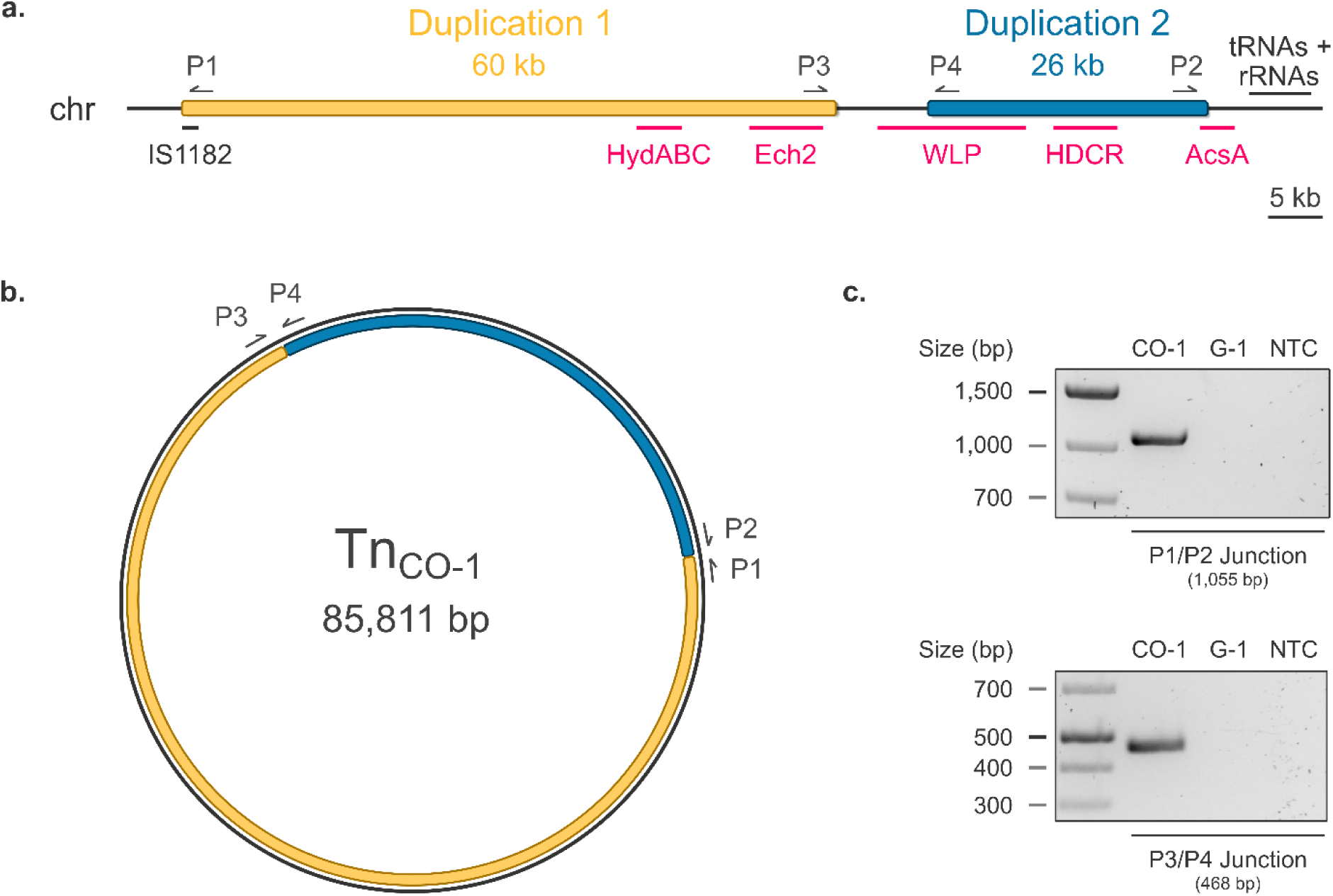
CO-1 large scale genome rearrangement identified by WGS. a) Coverage analysis (Illumina) highlights two regions duplicated in CO-1 encompassing most of the WLP locus as well as other gene clusters important for acetogenesis. b) Long read sequencing (ONT) reveals that both duplicated regions form a large, single, circular extrachromosomal element (Tn_CO-1_) containing an IS1182-like transposase gene. c) Validation of the presence of TnCO-1 in CO-1 DNA by PCR. Tn_CO-1_-specific junctions were amplified using P1/P2 and P3/P4 primers. NTC: no template control.

Tn_CO-1_ presence was next assessed in CO-2, CO-3, H-1 and H-2 strains by searching specifically for junction sequences, defined as 50 bp sequences linking the last 25 bp of each duplicated sequence. Tn_CO-1_ could be found in all strains able to grow on CO (CO-2, CO-3, H-1) but was absent from strains unable to grow on CO (H-2). On the other hand, *de novo* assembly of the H-2 genome showed that it bears the 2,438 bp deletion found in CO-1 (similarly to CO-2, CO-3, and H-1), indicating that this deletion probably does not play an important role in adaptation to CO.

### Tn_CO-1_ mobilization is CO-inducible

When changing the carbon source to glucose, we noticed long lag phases that were also observed when reverting from glucose to CO. We hypothesized this could be due to a dynamic regulation of Tn_CO-1_ favoring or impeding growth depending on the carbon/energy source. To test this hypothesis, we first performed semi-quantitative PCR targeting transposon junctions on DNA extracted from CO-1 and G-1 after a single culturing step on glucose, H_2_/CO_2_ and CO (the latter for CO-1 only, Figure 4a). Abundance of Tn_CO-1_-specific amplicons markedly increased when cells were grown on CO, indicating a higher prevalence of the megatransposon in this condition.

**Figure 4:**
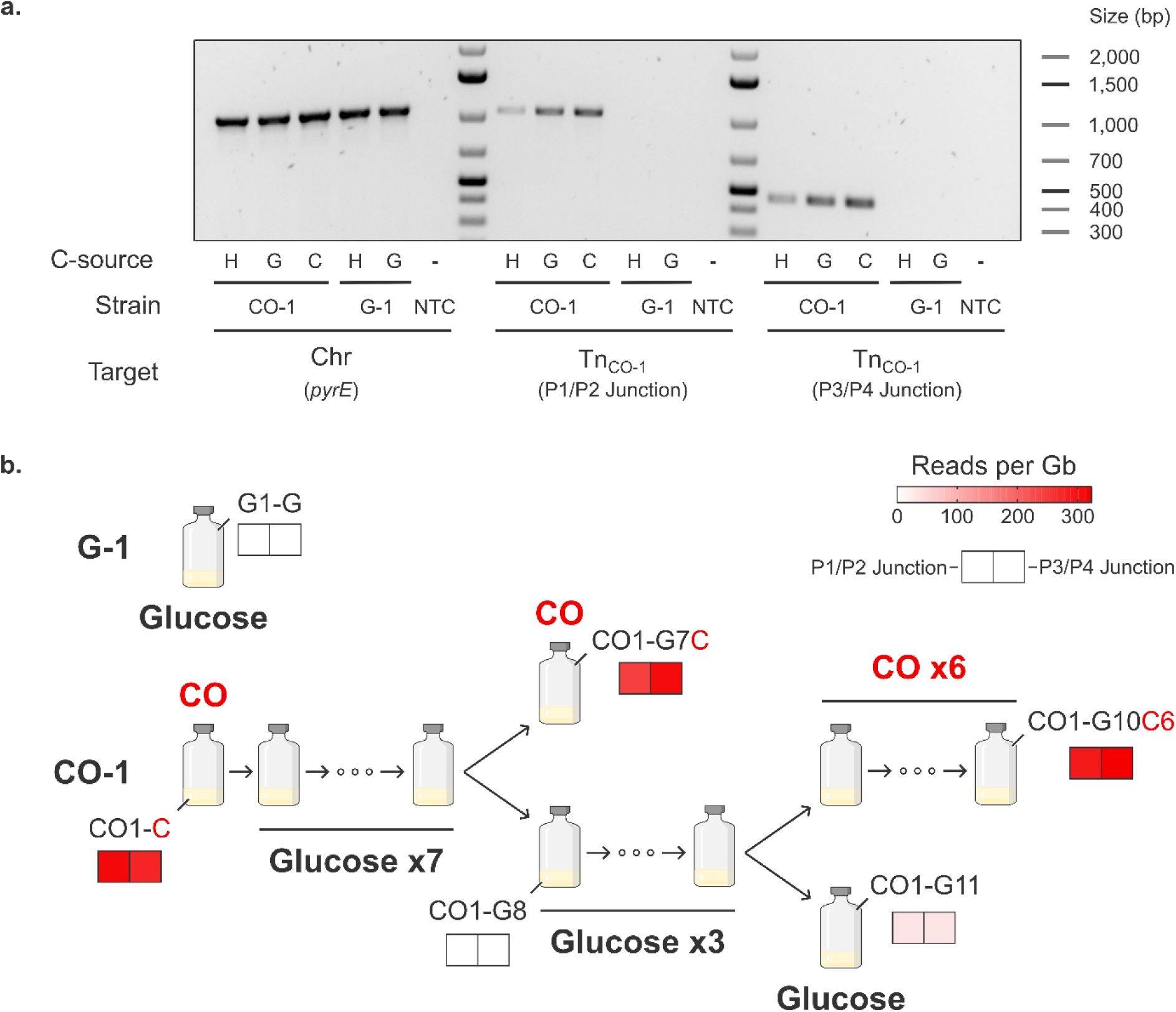
TnCO_-1_ mobilization is CO-inducible. a) Semi-quantitative end-point PCR was performed on gDNA extracted from CO-1 and G-1 strains grown under different carbon sources. H: 2 bar H_2_/CO_2_ (80:20). G: 5 g/L glucose. C: 2 bar CO. Shown are a PCR targeting chromosomal DNA (*pyrE* gene), as well as both Tn_CO-1_-specific junctions. NTC: no template control. b) Adaptation to glucose and readaptation to CO was monitored using deep sequencing data (ONT). Tn_CO-1_ presence was assessed by quantifying the abundance of Tn_CO-1_-specific junction sequences.

Tn_CO-1_ is extrachromosomal, and a decrease in abundance over time on alternative carbon sources could potentially lead to a complete loss of carboxydotrophy. We therefore subcultured CO-1 repeatedly on glucose and assessed whether the strain was able to grow back on CO after seven or ten culturing steps (Figure 4b). Readaptation to CO was successful, and cells at various stages of the lineage were sequenced using ONT. Tn_CO-1_ abundance was quantified and shown to be at best very low when cells were grown on glucose (maximum 34 reads per Gb, compared to ∼300 on CO). In contrast, re-exposure to CO resulted in much higher levels of Tn_CO-1_-specific reads, even after prolonged culturing on glucose. In this experiment, mobilization of Tn_CO-1_ was therefore strongly CO-inducible, which provides additional evidence of its importance for carboxydotrophy.

### Carboxydotrophic growth is abolished in the absence of Tn_CO-1_

Wild-type *T. kivui* is naturally competent^27^, which makes the strain genetically tractable. However, our initial attempts to transform the CO-1 strain with an empty *Thermoanaerobacter* – *E. coli* shuttle vector were unsuccessful. We therefore adapted our transformation protocol^28^ based on the observation that 55 °C as growth temperature gave higher transformation efficiency than 66 °C in the WT (data not shown). Low plating efficiency was circumvented by selecting neomycin-resistant strains in liquid before plating. Using this approach, we managed to isolate a CO-1 transformant bearing an empty backbone (Figure 5a). Surprisingly, the resulting strain was no longer able to grow on CO and lost most of its capacity to grow on high CO syngas (H_2_/CO_2_/CO: 25/25/50, Figure 5b). DNA was extracted from cells grown on low CO syngas (H_2_/CO_2_/CO: 60/15/25, a mixture also allowing low growth of the WT) and genotyping further showed that transformation of CO-1 resulted in the loss of Tn_CO-1_ (Figure 5c). This was confirmed by long-read sequencing, which failed to detect any reads indicative of the presence of Tn_CO-1_, while confirming all CO-1-specific mutations to be present. Taken together, our results therefore unambiguously show that the highly efficient carboxydotrophic lifestyle displayed by CO-1 is solely linked to the mobilization of the megatransposon Tn_CO-1_.

**Figure 5:**
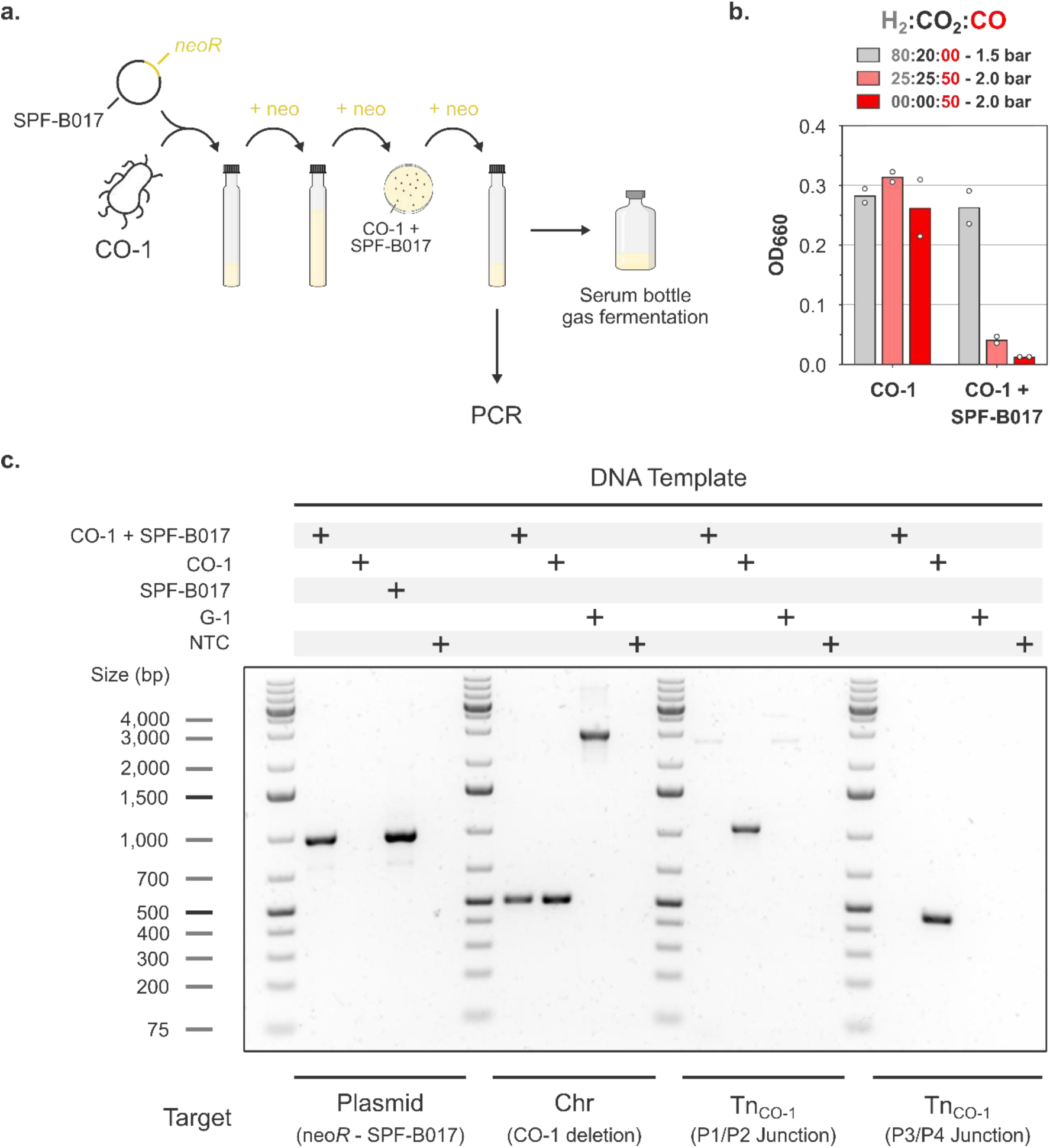
Transformation of CO-1 results in loss of carboxydotrophy. a) Scheme depicting a transformation procedure adapted to the low transformation efficiency of CO-1. CO-1 was transformed with the empty SPF-B017 shuttle plasmid. b) Biomass formation of non-transformed and transformed CO-1 after 3 days in serum bottles (mineral medium) under various gas mixtures. c) PCR assay on DNA extracted from non-transformed and transformed CO-1. Shown are PCR targeting SPF-B017, a CO-1-specific deletion, as well as both Tn_CO-1_-specific junctions. NTC: no template control.

### The CO-1 strain can efficiently convert CO-containing gasses in chemostat cultures

To characterize the physiology and study gene expression levels of strain CO-1, we established chemostat gas fermentations. This system provides well-defined, quantitative, stable steady-state conditions, and enables precise control of the specific growth rate in bioreactors. Triplicate continuous cultures of G-1 and CO-1 were set up in 0.25 L bioreactor vessels with continuous gassing, adjusting stirring speed to reach about 2 g L^−1^ acetate in the fermentation broth. G-1 and CO-1 were both cultivated on H_2_/CO_2_, CO-1 additionally on CO as sole carbon and energy source at two different specific growth rates.

Specific gas consumption and acetate production rates, as well as biomass and acetate yields were quantified for all conditions (Table 2). On H_2_/CO_2_ with a dilution rate D of 0.10 h^−1^, acetate production yields were similar between CO-1 and G-1, with about 4 H_2_ and 2 CO_2_ consumed per acetate formed, as expected from typical acetogenesis^29^. On CO (D = 0.10 h^−1^), high specific CO consumption rates were associated with high biomass yields, which might be linked to higher ATP yields from the carboxydotrophic lifestyle^10^. As a result, the acetate yield was higher than the theoretical yield (∼5.5 instead of 4 CO per acetate^30^). Similar to batch cultivations, CO-1 produced H_2_ and CO_2_ through the water gas shift reaction in chemostats.

**Table 2:**
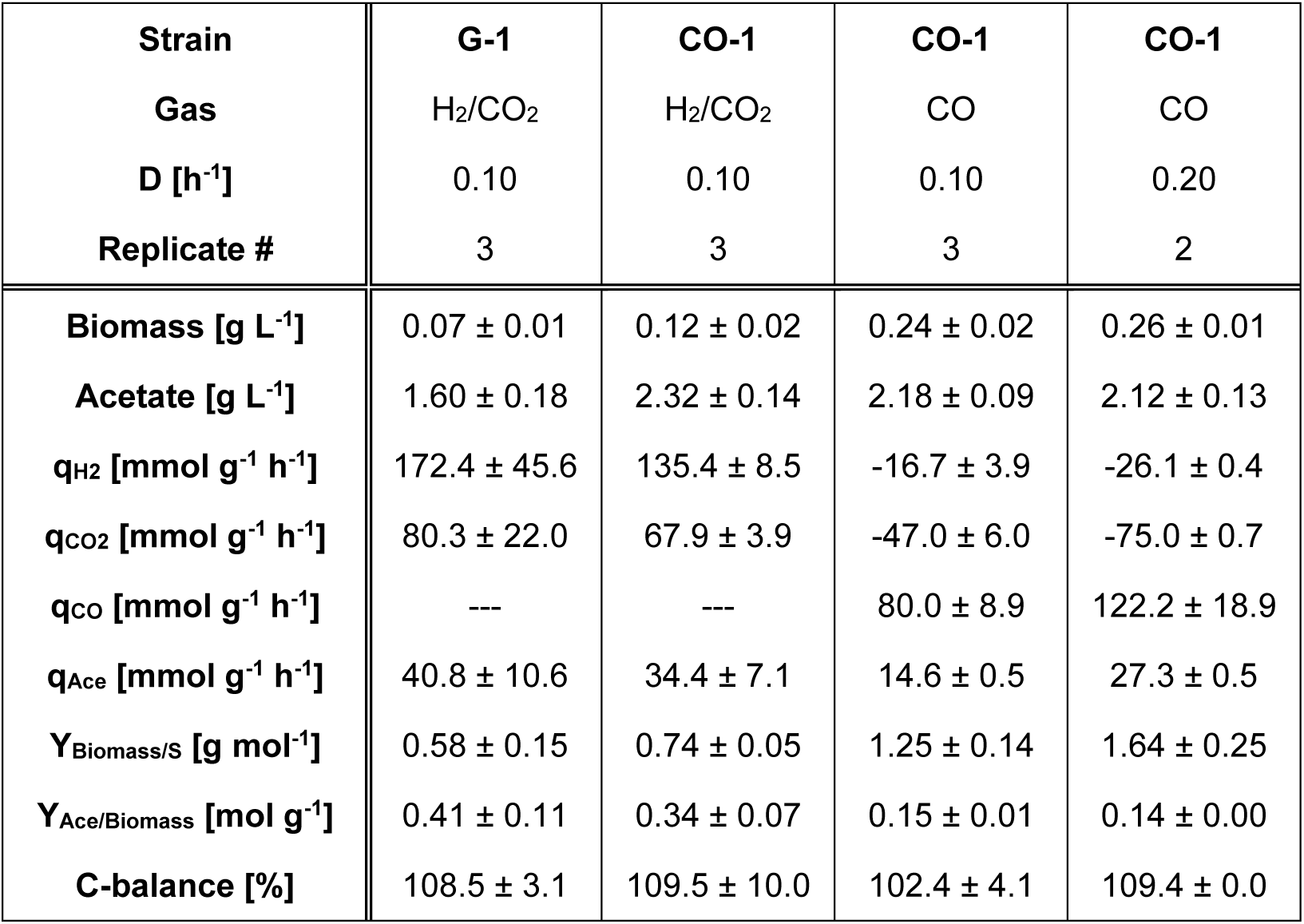
Fermentation metrics of steady-state continuous fermentation of G-1 and CO-1 grown in mineral medium using H2/CO2 or CO as sole carbon and energy sources.

Next, we established cultures at D = 0.20 h^−1^ on CO to compare the behavior of CO-1 at two different growth speeds. The gas-liquid mass transfer rate was adjusted by increasing the stirrer speed to maintain biomass and acetate titers. Compared to D = 0.10 h^−1^, a higher specific CO uptake rate (122.2 ± 18.9 mmol g^−1^ h^−1^) and a higher biomass yield (1.64 g mmol^−1^) was observed, showcasing the potential of strain CO-1 for bioprocessing. Finally, increasing dilution rate in a dynamic shift experiment until 0.25 h^−1^ was possible, thereby quantifying the maximum growth rate of the strain on CO.

### Steady-state transcriptomics analysis illuminates metabolic changes enabling carboxydotrophic growth

Samples for transcriptomic analyses were collected from steady-state cultures at D = 0.10 h^−1^ (Table 2). Gene expression was quantified, and differential expression analysis was used to gain insights between strains and carbon/energy sources (Supplementary Table S3). Principal component analysis indicated that the choice of the strain had the strongest impact on expression variance (Supplementary File S2). Comparison of the G-1 (H_2_/CO_2_) and the CO-1 (H_2_/CO_2_ or CO) datasets showed sporulation genes to be highly upregulated in CO-1 (19 genes with log_2_(FC) > 1 and adj. p-value < 0.05), suggesting that CO-1 might highjack part of the sporulation signaling cascade to cope with CO despite the inability of *T. kivui* to form mature spores^31^.

The strong carboxydotrophic phenotype exhibited by CO-1 could very well be the result of expression changes directly affecting the WLP, as most of the corresponding genes are born by the megatransposon Tn_CO-1_. Therefore, we investigated the strain-specific regulation of genes involved in CO metabolism (Figure 6). In CO-1, transcriptomics data indicated a potential reorganization of acetogen metabolism, with expression levels of many redox-related genes being significantly regulated. The NAD^+^-dependent electron bifurcating NADPH:Fd oxidoreductase Nfn, and the Ech2 complex were significantly upregulated, with Ech2 exhibiting the highest changes (log_2_(FC) between 1.33 and 2.52). Ech2 as well as Nfn both use reduced ferredoxin (Fd^2-^) suggesting that the cellular balance between reduced and oxidized ferredoxin (Fd) might be critical for carboxydotrophy.

**Figure 6:**
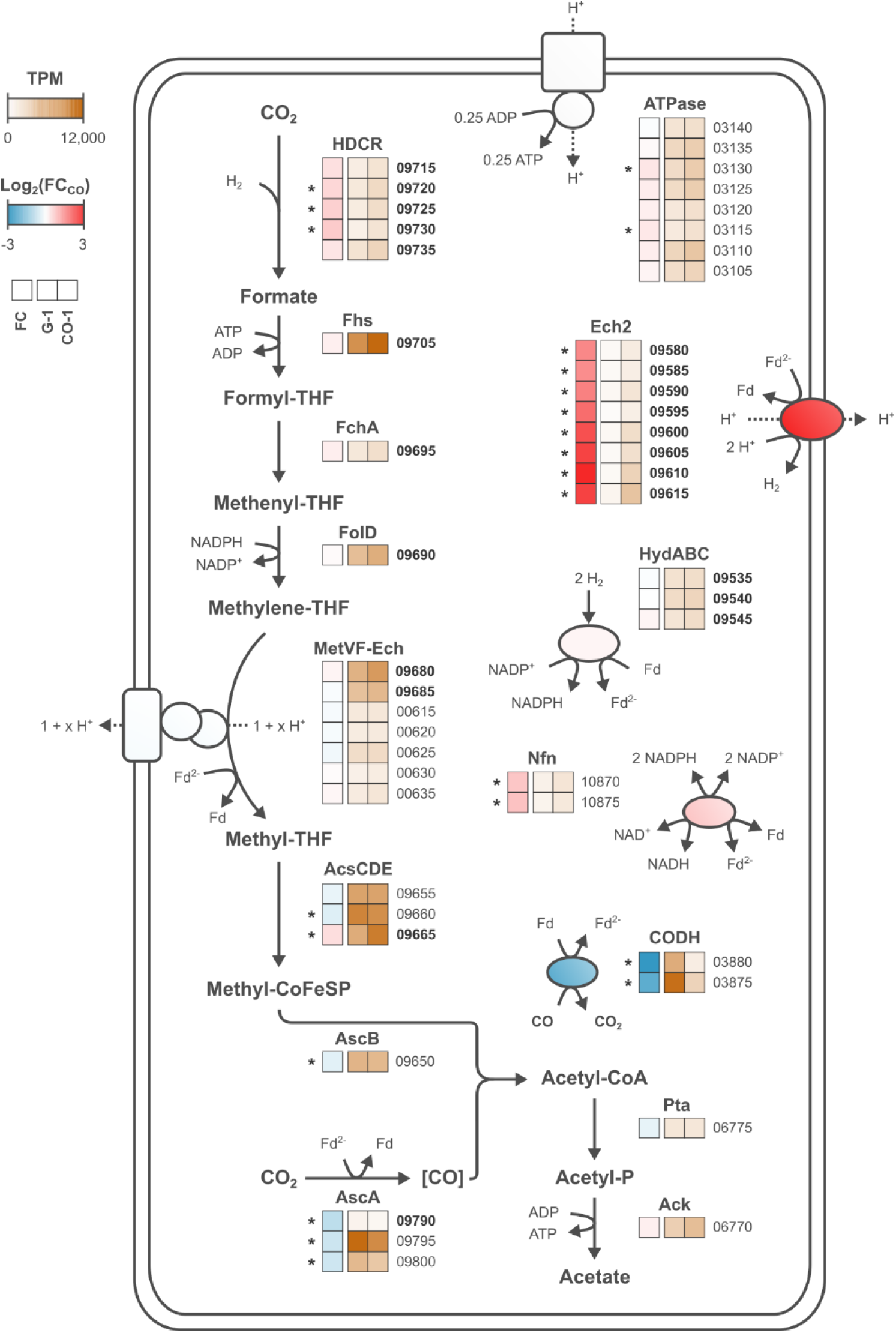
Steady-state transcriptomics view of metabolism in G-1 and CO-1 strains. Central metabolism comprising the WLP as well as genes involved in energy conservation are shown. Mean TPM values as well as fold changes for G-1 (H_2_/CO_2_) and CO-1 (CO) are described as a heat map. Locus tags in bold indicate genes present on Tn_CO-1_. Asterisks correspond to statistically significant values (adjusted p-value < 0.05). RNA-seq was performed using RNA extracted from triplicate continuous fermentation experiments described in Table 2.

The monofunctional CO dehydrogenase CODH genes were counter-intuitively downregulated in CO-1 (log_2_(FC) of –1.96 and –2.45 for *cooS* and *cooF1*). Indeed, CODH is essential for CO metabolism^32^. Similarly, the acetyl-CoA synthase Acs genes were also slightly repressed, although CODH and Acs are the only enzymes directly utilizing CO. Similar to Ech2 and Nfn, both enzymes are linked to the cellular Fd^2-^/Fd pool. Reduced expression of CODH, an enzyme with high turnover rates^32^, is expected to strongly limit CO oxidation, and therefore the overgeneration of Fd^2-^. This synergizes with increased Ech2 activity, as Ech2 consumes Fd^2-^. In this view, maximizing regeneration of Fd would be linked with the carboxydotrophic phenotype, in a similar fashion that maximizing NAD^+^ regeneration favors efficient glucose uptake in glycolytic microorganisms.

### Overexpression of Ech2 is sufficient to promote carboxydotrophic growth in wild-type *T. kivui*

In our transcriptomics dataset, Ech2 upregulation in the CO-1 strain was particularly strong, and in the context of its inhibition by CO and its putative role in balancing the Fd pool, we hypothesized that the Ech2 expression level might be crucial for CO adaptation. The mechanism underlaying the upregulation of Ech2 in the CO-1 strain is likely linked with a reorganization of the WLP genes on Tn_CO-1_ (Figure 7a). Indeed, the strongly expressed *acs* genes are interrupted at the level of *acsC* (TKV_RS09660) in Tn_CO-1_, thereby removing the putative terminator located downstream of *gcvH* (TKV_RS09645). This regulatory element (identified with ARNold^33^) should terminate transcription of the entire WLP locus (organized in a single operon starting with *fhs*, TKV_RS09705, based on transcriptomic data). Consequently, in Tn_CO-1_ the *ech2* genes are likely both under the control of their native promoter as well as the significantly stronger WLP promoter (about 11.5-fold stronger based on G-1 transcriptomic data).

**Figure 7:**
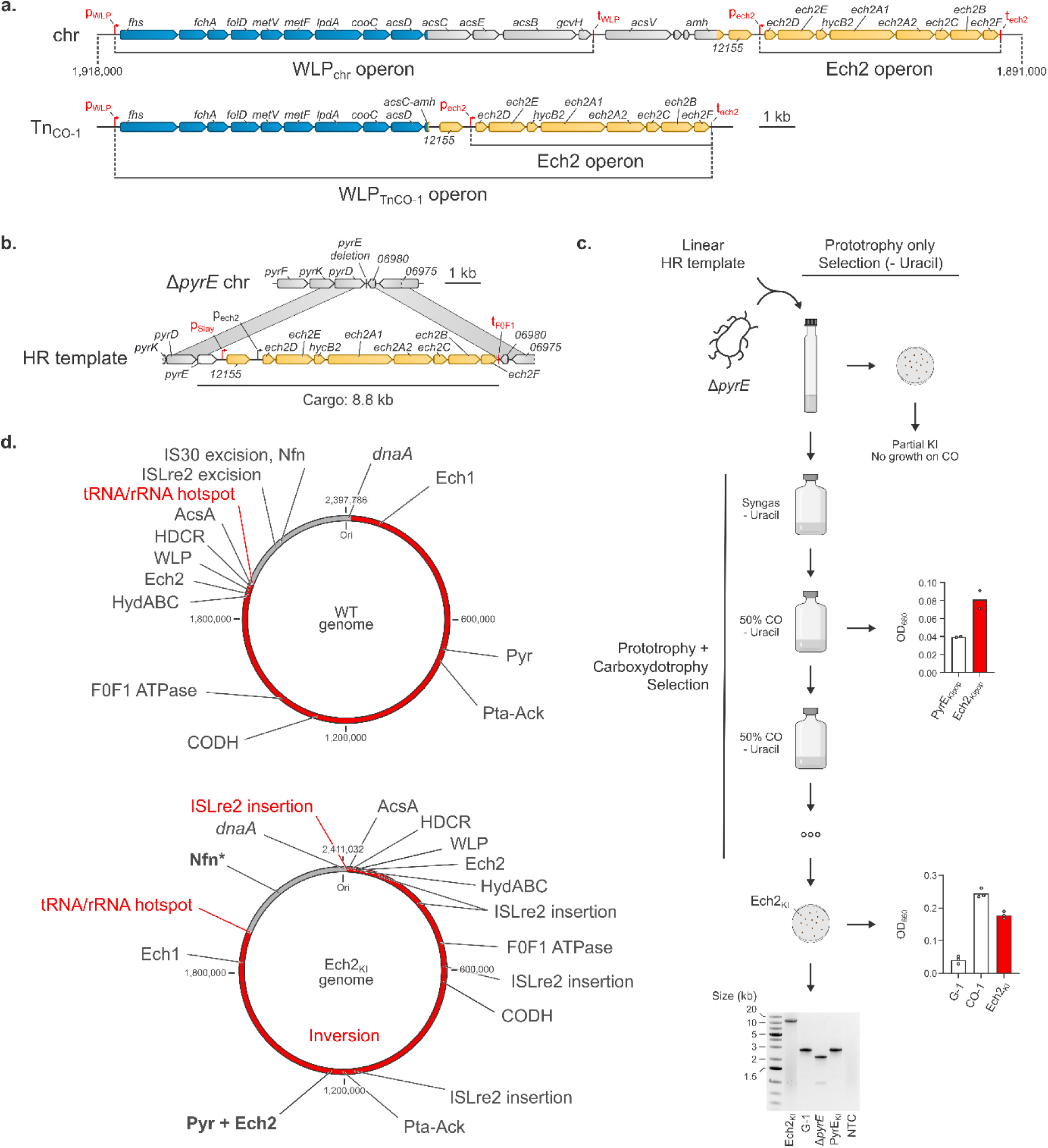
The Ech2 enzyme is crucial for CO adaptation in *T. kivui*. a) Overview of the WLP and Ech2 locus organization on the chromosome and Tn_CO-1_. b) Genome editing design for the overexpression of Ech2. c) Genome editing pipeline: direct selection of prototrophic cells, versus coupled selection of prototrophy and carboxydotrophy. NTC: no template control. d) Large– and medium-scale rearrangements in the carboxydotrophic Ech2_KI_ strain. WT refers to GCA_963971585.1. Bold: mutated genes. Except for *dnaA*, features refer to loci and not genes.

We therefore sought to replicate a genetic organization similar to Tn_CO-1_ by knocking in all *ech2* genes (TKV_ RS09580-9615), as well as the hypothetical CDS upstream of these genes (TKV_RS12155, also significantly upregulated, log_2_(FC) = 2.50), in a wild-type background. We chose the strong, constitutive S-layer promoter^28^, and a gene editing system based on the selective/counter-selective auxotrophic marker *pyrE* (conferring uracil prototrophy) and linear DNA transformation as previously described^27^ (Figure 7b). A 10.8 kb PCR fragment designed to knock in *pyrE* and *ech2* genes at the *pyrE* locus was used to transform a Δ*pyrE* strain derived from the wild type (Figure 7c). Selection without uracil coupled to random isolation on plates yielded multiple strains displaying correct integration of *pyrE* but loss of all or most of the cargo (data not shown). The same cultures used for isolation were subcultured, using 50 % CO as a selective pressure. After prolonged incubation at 66 °C, an increase of OD_660_ was observed for the Ech2 knock-in transformations, but not for the negative control (knock-in of *pyrE* without any additional cargo). Further subculturing showed the ability of the culture to regrow on CO as sole carbon and energy source, and individual clones were isolated on plates and screened for carboxydotrophy. A single isolate, referred to as Ech2_KI_, was sequenced with both long– and short-read sequencing, which revealed correct integration of the cargo at the *pyrE* locus, in contrast to the direct isolation step. This indicates that correct integration of the Ech2 knock-in was likely selected for by CO exposure, further demonstrating the importance of Ech2 for carboxydotrophic growth.

### The Ech2_KI_ strain bears additional small-to large-scale mutations

Since the selection step on 50 % CO required one month, we checked for other mutations in the Ech2_KI_ strain that could have contributed to establish robust carboxydotrophic growth. In particular, we hypothesized that strong overexpression of the membrane-bound Ech2 complex could impose a metabolic burden, and that the expression level of Ech2 subunit genes might therefore require refining.

SNV/InDel analysis identified 21 mutations (Supplementary Table S4). A handful of the newly acquired mutations could play a significant role in rewiring acetogen metabolism, including pleiotropic regulators such as *codY* or *phoU*, both of which could modulate carbon metabolism^34,35^. The Nfn subunit NfnB was also significantly extended in Ech2_KI_ (463 AA instead of 407). Importantly, the C-terminus of NfnB is involved in NADPH binding^36^, and the extension might therefore affect the redox activity of the whole complex.

In addition to SNV/InDel analysis, we performed *de novo* assembly of the Ech2_KI_ genome and found significant medium– and large-scale rearrangements, extensively remodeling the genomic landscape (Supplementary Table S4, Figure 7d). High transposon activity resulted in the insertion or excision at multiple loci of single gene transposases, locally affecting several genes. Most of this transposon activity was mediated by an ISLre2 family transposase, i.e., a different transposase as the one involved in formation of the megatransposon Tn_CO-1_ in CO-1. Transposon excision notably occurred in close proximity to the Nfn encoding genes (upstream of TKV_RS10865, encoding a secondary alcohol dehydrogenase), which might affect *nfnAB* expression. Whole genome alignment further revealed the inversion of a 1,937 kbp fragment (approximately 80 % of the wild-type genome). This inversion is flanked on one side by an ISLre2 family transposase, and on the other side by the same tRNA/rRNA hotspot identified at the end of the CO-1 duplication (Figure 1a). The inversion completely reshapes the genome (Figure 7d), and notably 1) brings the WLP locus much closer to the origin of replication (at about 15 kbp from the start) and 2) pushes the knock-in locus away from the origin of replication (at about 1,150 kbp, halfway through the genome). In bacteria, the position of a gene relative to the origin of replication strongly affects its expression, with genes closer to the origin of replication being on average more expressed^37^. In Ech2_KI_, the novel positional bias resulting from the inversion is therefore expected to increase the expression of all genes from the WLP locus, while decreasing the expression of the Ech2 knock-in. Stronger expression of the WLP locus could be selected for on CO as a way to increase carbon flux through the WLP, and/or to increase the availability of enzymes typically inhibited by CO (e.g., the HDCR). Simultaneously, a lower level of overexpression of Ech2 could decrease the metabolic burden linked with high expression, optimizing Ech2 levels to accommodate carboxydotrophic growth.

## Discussion

In this work, we adapted *T. kivui* to grow solely on CO, and rapidly obtained strain CO-1 within 31 generations which tolerates and uses CO efficiently. Growth on CO was shown to be particularly fast (µ_max_ = 0.25 h^−1^, T_d_ ∼ 2.8 h), in particular compared to values reported previously for CO-adapted *T. kivui* (0.017 h^−1^, T_d_ ∼ 40.8 h)^7^. To the best of our knowledge, the growth rate of CO-1 is higher than the growth rate of any other acetogen grown on CO in chemically defined medium. The closest µ_max_ was reported for *C. autoethanogenum* in chemostat cultivations (0.12 h^−1^, T_d_ ∼ 5.8 h)^38^, highlighting the relevance of our phenotype for functional analysis and for bioprocessing of syngas. We further showed that the strong carboxydotrophic ability of CO-1 is conferred by the mobilization of the CO-inducible extrachromosomal megatransposon Tn_CO-1_. Indeed, this mobile element is essential to mediate the CO-1 phenotype, as Tn_CO-1_ deletion in CO-1 results in complete loss of carboxydotrophy.

How Tn_CO-1_ is mobilized and rearranges two genomic loci into a single circular DNA molecule however remains elusive, as little information is available for IS1182-like elements. Mobilization of a single 94 kb DNA fragment could occur with a copy-in-paste-out /circularization mechanism^39^, followed by excision of the 8 kb fragment at the WLP locus. Alternatively, the two genomic regions could be mobilized separately and subsequently fused, although this process is less probable since the IS1182-like transposase gene is only present once in Tn_CO-1_. Regardless of the activation and formation mechanisms, Tn_CO-1_ mobilization turned out to be remarkably stable in CO-1, as it could be consistently initiated by addition of CO.

In acetogens, inhibition of [FeFe]-hydrogenases is thought to be the primary driver of CO toxicity^10,13,17^. Here, we characterized an exceptionally robust carboxydotrophic acetogen, in which the major [FeFe]-hydrogenases HydABC and HDCR where neither mutated nor significantly up-or downregulated (Figure 6). In contrast, the Ech2 [NiFe]-hydrogenase was significantly upregulated. Ech2 was previously shown to be inhibited by CO^11^, however the IC_50_ value (∼ 200 µM, compared to ∼ 0.25 µM for the HDCR) suggests Ech2 inhibition does not occur *in vivo*, as the physiological intracellular CO concentrations are expected to be below the µM range^40^ and given that CODH activity was likely to be sufficient to completely oxidize CO diffusing across the cell membrane^41^. In addition, previous work showed that a Ech2 knock-out prevented adaptation of *T. kivui* to CO + yeast extract^17^, while our study demonstrated Ech2 KI could kickstart growth on CO as sole energy/carbon source. Ech2 provides a key activity as it allows the regeneration of Fd. As the Fd^2-^-generating CODH is also strongly downregulated in CO-1, the differential expression of both enzymes likely synergizes by limiting Fd^2-^ generation. In this metabolism-centered view of CO toxicity, the accumulation of Fd^2-^ through CO oxidation with the simultaneous depletion of the Fd pool is detrimental for central metabolic reactions (e.g., at the HydABC and Nfn levels), effectively paralyzing metabolism.

During the final stages of this study, Baum et al. (2024) published Illumina data of CO-adapted *T. kivui* strains^17^. SNP/InDel analysis suggested CO adaptation derived from a mutation arising in the HDCR complex. By performing coverage analysis, we uncovered an unreported ∼125 kb duplication in 4 out of 5 analyzed clones, between position ∼125 kb and 250 kb of the published genome of *T. kivui* DSM 2030 (NZ_CP009170)^42^. In this region, average coverage is ∼ 3.4-fold compared to the rest of the genome, and interestingly stops at a ISLre2 transposase gene (TKV_RS01200, the same involved in Ech2_KI_ inversion), suggesting that another megatransposon might be at play in CO adaptation of the corresponding clones. Altogether, many genes are duplicated, including some that might be highly relevant for carboxydotrophic growth, such as those encoding the energy-converting hydrogenase complex Ech1 subunits (TKV_RS00615-0655), and gene expression analysis showed upregulation of Ech1 subunit genes. However, the role of Ech1 in *T. kivui* metabolism is not fully understood and so far no Fd^2-^ oxidation activity has been determined for Ech1^17^. In contrast, upregulation of Ech2 genes also found in these strains might suggest an underlying CO tolerance mechanism at least partly similar to the one in CO-1, where balancing the Fd^2-^/Fd pool appears to be crucial.

Efficient consumption of excess Fd^2-^ by Ech2 under highly reduced CO fermentation is therefore likely the key to promote carboxydotrophy in *T. kivui*, provided their expression is sufficient to balance CODH-mediated ferredoxin reduction. Indeed, Ech2-mediated Fd^2-^ regeneration can serve as a “redox pressure relief valve” yielding H_2_, which can escape the cell (which is supported by our experiments, as we observed H_2_ production from CO). Additionally, carbon reduction in the methyl branch of the WLP could be an alternative route for Fd^2-^ reoxidation as Fd^2-^-dependent methylene-THF reductase activity has been demonstrated^30,44^.

In the natural carboxydotrophic acetogen *Clostridium autoethanogenum*, a CO tolerance mechanism in principle similar to the one we describe for *T. kivui* has been proposed. Indeed, the ethanol fermentation pathway via the aldehyde:ferredoxin oxidoreductase (AOR) of *C. autoethanogenum* plays a crucial role to maintain a balance between Fd and Fd^2-^ by consuming Fd^2-^ under high CO concentration, using acetate as an electron sink^41,46–48^. In *Eubacterium limosum*, adaptation to CO resulted in mutations affecting the CODH subunit of the acetyl-CoA synthase (AcsA or cognate maturation protein CooC2)^8^, which, in the absence of a monofunctional CODH is also responsible for CO oxidation. Although the mutations were not reverse engineered, it is highly possible that these mutations negatively affect AcsA-mediated Fd reduction.

As a result, adaptation of *E. limosum* to CO could follow the same rules as in *T. kivui* CO-1. In *A. woodii*, recent results have shown that mixotrophy on CO + yeast extract is possible when the HDCR is engineered from an H_2_-consuming enzyme to a Fd^2-^-consuming enzyme^45^, suggesting fine-tuning the redox balance could also be at play. In that case, circumventing the CO inhibition occurring at the HDCR level might be equally or more important, as the Fd^2-^ activity of the HDCR is not CO-inhibited^12^.

Thermophiles in general and *Thermoanaerobacter* species in particular tend to display low genetic barriers with, e.g., highly efficient natural competency^49^, favoring in nature horizontal gene transfer (HGT). Thermophilic bacteria live in “extreme” environments, which requires fast adaptive capability^50^. Interestingly, *T. kivui* is the only *Thermoanaerobacter* described so far capable of autotrophic growth. Because most genes involved in the WLP are colocalizing in the *T. kivui* chromosome^42^, acquisition of these genes was hypothesized to stem from a single large HGT event^18^, suggesting that autotrophy could be fully transferrable through horizontal exchange. That view was recently challenged, as a phylogenetic analysis suggested autotrophy was vertically inherited, and lost in all *Thermoanaerobacter* sp. but *T. kivui*^51^. Throughout our study, we witnessed two cases of mobility of the WLP locus: first through the mobilization of Tn_CO-1_, and second via a large inversion of the genome. Both events were likely mediated by transposases (from the IS1182, and ISLre2 families, respectively), as these were found in close proximity of the corresponding rearrangements. In both cases, a tRNA/rRNA hotspot (TKV_RS09805-9855) marks the other end of the rearrangement. tRNA and rRNA genes are strategic targets for transposition elements, as their sequences are highly conserved among taxa, favoring cross-species transfer of these mobile elements^21–26^. Collectively, the high mobility of the DNA segment bearing the WLP genes, the involvement of transposases and the presence of a tRNA/rRNA hotspot all support the hypothesis that *T. kivui* acquired autotrophy via horizontal gene transfer, adding yet another example of the fundamental role transposons have played and keep on playing in the evolution of life^52,53^.

## Online Methods

### Strains, medium and culture conditions

Supplementary Table S5. contains the strains used in this study. The strains were cultivated on mineral medium at 66 °C or 55 °C as previously described^28^. Glucose (5 g L^−1^) or gas was used as carbon/energy source as necessary in serum bottles or Hungate tubes sealed with rubber stoppers. For glucose fermentation, 2 bar N_2_/CO_2_ (80:20) was used as make-up gas. For gas fermentation, the headspace of serum bottles was flushed with the appropriate gas, and the pressure was set to 2 bars, unless otherwise stated. Gasses for the serum bottles were mixed with Brooks 4800 series mass flow controllers (Brooks Instrument, Hatfield, USA), except for CO, which was added separately and the concentration set by fraction of pressure. Medium was supplemented with 2 g L^−1^ yeast extract (for cultivating the Δ*pyrE* strain only), and with 200 mg L^−1^ kanamycin or 100 mg L^−1^ 5-fluoroorotic acid when appropriate.

For solid medium, mineral medium was supplemented with 7 g L^−1^ gelrite, 5 g L^−1^ glucose and 2 g L^−1^ yeast extract (except for uracil prototrophy selection). 200 mg L^−1^ neomycin was added when appropriate. Cells were grown in custom-made metallic jars pressurized at 2 bar N_2_/CO_2_ (80:20) at 55 °C or 66 °C until colony formation.

For bioreactor experiments, the batch medium was modified to remove carbonate and to contain an additional 0.750 g L^−1^ NH_4_Cl. For chemostat, the batch medium was also supplemented with 28.6 mg L^−1^ FeSO_4_, 7 H_2_O.

*E. coli* DH10B (Thermo Fisher, MA, USA) or CopyCutter EPI400 (Epicentre, WI, USA) were cultivated on LB medium (10 g L^−1^ tryptone, 10 g L^−1^ NaCl, 5 g L^−1^ yeast extract) with 50 mg L^−1^ kanamycin, 100 mg L^−1^ carbenicillin and 1X CopyCutter induction solution when appropriate. For solid medium, 15 g L^−1^ agar was added.

### Adaptation to CO

*T. kivui* DSM 2030 was obtained from the DSMZ collection and directly used for CO adaptation without reisolation. CO adaptation was undertaken by serially growing *T. kivui* in 125 mL serum bottles filled with 20 mL mineral medium and 2 bars of the appropriate gas. Cells were grown at 66 °C with shaking in a water bath (Memmert, Schwabach, Germany). Upon visible growth, 1 mL of cell culture was transferred to the next serum bottle for every adaptation step, until high growth on 100 % CO was observed. For clonal strain isolation, cells were streaked on plates and isolated colonies were grown in Hungate tubes with 2 bars of the appropriate gas mixture.

### Fermentation in bioreactor

Bioreactor cultivations were carried out in a DASBox® Mini Bioreactor system (Eppendorf AG, Hamburg, Germany). Batch cultivations were conducted with a working volume of 250 mL and an agitation rate of 600 rpm for the growth on syngas (H_2_/CO_2_/CO/N_2_: 24/21/52/3) and an agitation of 300 rpm for the growth on 100% CO. An aeration rate of 0.05 vvm was applied.

Continuous cultivations were carried out with a working volume of 208 mL. Agitation was adjusted to maintain a constant acetate titer for the growth on H_2_/CO_2_ (H_2_/CO_2_/N_2_: 58/9/33) and CO (CO/N_2_: 52/48) with an aeration rate of 0.06 vvm. All gas mixtures were premixed and obtained from Messer (Messer Austria GmbH, Gumpoldskirchen, Austria).

All bioreactor cultivations were operated at a constant temperature of 66 °C using a static water bath. The pH was set to 6.4 and constantly monitored with an Easy Ferm Plus K8/120 pH electrode (Hamilton, Reno, NV, USA) and automatically adjusted by the addition of 5 M KOH using a MP8 multi pump module (Eppendorf AG, Hamburg, Germany). Uniform gassing was ensured using a sintered microsparger with a defined pore size of 10 µm (Sartorius Stedim Biotech GmbH, Göttingen, Germany). Prior to inoculation the reactor was flushed with the appropriate gas mixture for at least 3 h to ensure anaerobic conditions. The bioreactors were inoculated to an initial OD_600_ of 0.1. Chemostat cultivations were switched to continuous mode after the pH regulation was activated for the first time.

Reactor sampling was carried out every 2.5 h in batch mode and once per day for continuous mode, monitoring OD_600_ (ONDA V-10 Plus Visible Spectrophotometer) and product content by HPLC (Ultimate 3000 High Performance Liquid Chromatograph, Thermo Fisher Scientific, Waltham/MA, USA). For HPLC, an Aminex HPX-87H column (300 × 7.8 mm, Bio-Rad, Hercules/CA, USA) was used to analyze the samples, using a refractive index (Refractomax 520, Thermo Fisher Scientific, Waltham, MA, USA) and a diode array detector (Ultimate 3000, Thermo Fisher Scientific, Waltham, MA, USA) for quantification.

For dry cell weight determination 50 mL of culture broth was collected at the end of the batch cultivation and at steady state condition for the continuous cultivation. The tubes were centrifuged and washed two times with 0.9 % (w/v) NaCl. The biomass sample was then suspended with 5 mL distilled water, transferred into pre-weighed glass tubes, dried out for 24 h at 120 °C and cooled down in a desiccator for 1 h before weighing. Biomass determination was performed in triplicates.

A Micro GC Fusion® Gas Analyzer (Inficon Holding AG, Bad Ragaz, Switzerland) was used to analyze the off-gas of all reactors every hour using a Valco stream selector from VICI® (Valco® Instruments Co. Inc., Houston, Texas). To account for the volume change in the off-gas, the N_2_ concentration was used for normalization of the gas composition. A Rt-Molsieve 5A column using argon as carrier gas was used to detect N_2_, H_2_ and CO. A second Rt-Q-Bond was used to detect CO_2_ with helium as carrier gas. A thermal conductivity detector was used for quantitative analysis. Online GC values were used to calculate gas uptake rates as in Novak et al^54^.

### Short-read sequencing

For strain resequencing (CO-1, CO-2, CO-3), cell pellets were harvested from growing cultures, frozen at –80°C and sent to Microsynth GmbH (Balgach, Switzerland). Cells were subsequently lysed with lysozyme overnight in STET buffer (10 mM Tris-HCl, 1 mM EDTA, 100 mM NaCl and 5% Triton X-100) at 37°C, followed by proteinase K / RNase A treatment. DNA was isolated from the supernatant using the DNeasy kit (Qiagen) according to manufacturer’s instructions. Illumina’s DNA Prep tagmentation library preparation kit was used according to manufacturer’s recommendations to construct the libraries. Subsequently the Illumina NovaSeq platform with an SP flow cell and a 200 cycles kit were used to sequence the libraries. Paired-end reads which passed Illumina’s chastity filter were subject to de-multiplexing and trimming of Illumina adaptor residuals using Illumina’s bcl2fastq software version 2.20.0.422 (without further refinement or selection).

For polishing *de novo* assemblies, Illumina reads were obtained from the same DNA sequenced with ONT. Cells were treated with lysozyme in TE buffer, and subsequent DNA isolation was performed using the Quick-DNA Miniprep Plus kit (Zymo Research, Irvine, CA, USA) according to the manufacturer’s instructions. CO-1 DNA was sequenced by Microsynth GmbH as described above, except the TruSeq DNA library kit was used. Ech2KI DNA library was prepared by PlasmidSaurus (Eugene, OR, USA) using the seqWell ExpressPlex 96 library prep kit according to manufacturer’s instructions and was further sequenced on a NextSeq2000 sequencer (paired-end, 2x150 bp).

### SNV/InDel analysis

A wild-type genome was first generated by using wild-type Illumina reads (obtained from the culture used for ALE) to polish the G-1 genome (<u>OZ020628</u>), resulting in assembly GCA_963971585.1. In brief, Illumina reads were trimmed and filtered with trimmomatic^55^ (v0.38) with default parameters except for a minimum read length of 100 bps; the trimmed reads were used to polish the assembly twice with minimap2^56^ (v2.24-r1122) and HyPo^57^ (v1.0.3) with default parameters. Finally, the assembly was validated by remapping the trimmed reads and calling SNVs/Indels with CLC Genomic Workbench (v6.5.2; Qiagen GmbH, Hilden, Germany). Two remaining loci with unclear sequence were subjected to Sanger sequencing and the sequence manually curated accordingly. For the analysis of the evolved strains, Illumina reads were mapped onto the wild-type assembly with Geneious Prime 2024.0.5 (https://www.geneious.com) using the minimap2^56^ plugin (v2.24, short read mode, default parameters). SNV/InDel analysis was performed with the built-in function of Geneious, filtering for mutations arising in more than 80% of the mapped reads for a given position.

### Long-read sequencing

Cell pellets were harvested from growing cultures, pelleted and directly processed for DNA extraction. Cells were treated with lysozyme in TE buffer, and subsequent DNA isolation was performed using the Quick-DNA Miniprep Plus kit (Zymo Research, Irvine, CA, USA) according to the manufacturer’s instructions. Extracted DNA was sent to PlasmidSaurus for ONT sequencing. Libraries were prepared with the Rapid Barcoding Kit 96 V14 (SQK-RBK114.96, ONT) according to the manufacturer’s instructions, without DNA shearing or size selection, before subsequent sequencing (PromethION, R10.4.1 flowcell). Raw reads were obtained after basecalling, barcode splitting and adapter trimming (Guppy v.6.4.6, super-high accuracy mode).

### *De novo* assembly

CO-1 strain: the 390,315 raw reads (N_50_ = 5,697) were trimmed with PoreChop^58^ v0.2.4, down-sampled to 100x coverage with Seqkit v2.4.0^59^ and sorted for longest reads, resulting in 15,957 reads (N_50_ = 15,344). Draft assembly was performed with Canu v2.2^60^ with default parameters, yielding 5 linear contigs. Based on the similarity to the DSM2030 genome, the longest contig (2,424,656 bp) was identified as the CO-1 chromosome. Overlaps of both ends of this contig were determined by BLASTn^61^, the chromosome circularized based on this overlaps and rotated to start at *dnaA* gene with Seqkit v2.4.0^59^. The genome was polished and validated using the corresponding Illumina data as described above, resulting in a 2,396,992 bp chromosome. Annotations were transferred from *T. kivui* G1 genome (GCA_963971585.1) with Geneious. For the 4 remaining contigs (36,204 bp, 35,793 bp, 55,987 bp and 60,479 bp), regions overlapping the chromosomal contig were found at the duplication sites previously determined through Illumina sequencing using BLASTn^61^. These 4 contigs significantly overlapped each other, which allowed the assembly of a single, circular extrachromosomal element (85,811 bp).

Ech2_KI_ strain: *de novo* assembly and polishing were performed by PlasmidSaurus. The 135,120 reads (N_50_ = 4,200) were filtered using Filtlong^62^ (v0.2.1, default parameters) to exclude the 5% reads with the lowest average quality. Reads were down-sampled to 100 x coverage with Filtlong, weighted by quality score. *De novo* assembly was performed with Flye^63^ (v2.9.1), with parameters selected for high quality ONT reads. The resulting assembly was polished using the down-sampled ONT reads with Medaka^64^ v1.8.0 and all Illumina reads with Polypolish^64^ v0.6.0, yielding a 2,411,034 bp chromosome, which was further annotated with Bakta^65^. Finally, the chromosome was rotated to start at *dnaA*.

### Structural variation analysis

CO-1 and Ech2_KI_ chromosomes were aligned to the wild type assembly using Mauve^66^. Transposon translocations and sequence inversion were manually inspected using Geneious Prime.

### Transposon quantification

Seqkit v2.4.0^59^ was used to select raw ONT reads bearing Tn_CO-1_-specific junctions sequences, defined as 50 bp sequences composed of 25 bp specific of each side flanking a given junction. P1/P2 junction sequence was 5’-aaaacagggcacttaagtgccctgttcagactgttgacaaaatatcggga-3’, and P3/P4 junction was 5’-agtatatcttttttatattatccatattgcaaaagcaagacaagttggaa-3’. For quantification, the number of reads bearing each junction was normalized using sequencing depth (in Gb).

### Genotyping

PCR was routinely used to assess genotype, by targeting chromosomal loci (*pyrE*, CO-1 specific deletion), SPF-B017 or Tn_CO-1_ (at P1/P2 and P3/P4 junctions). S7 Fusion polymerase (Biozym, Hessisch Oldendorf, Germany) was used to amplify DNA from purified genomic DNA or directly from cell cultures. For semi-quantitative PCR, gDNA was quantified with Qubit (Thermo Fisher), diluted to 2 ng µL^−1^ and used for PCR, limiting cycling to visualize DNA without saturation after gel electrophoresis.

### Transcriptomics

Cells from steady-state chemostats were pellets and snap-frozen in liquid nitrogen. Pellets were next shipped on dry ice to Microsynth GmbH for processing. DNA/RNA Shield (Zymo Research, Irvine, CA, USA) was used to resuspend cell pellets, and cells were lysed by bead beating. Soluble fraction was thereafter used for RNA isolation using the RNeasy Plus 96 Kit (Qiagen). Libraries were prepared with the Stranded Total RNA Prep Kit with Ribo-Zero Plus Kit (Illumina), following the manufacturer’s instructions. Sequencing was performed on an Illumina NovaSeq 6000 device (2 x 150 bp). The paired-end reads, which passed Illumina’s chastity filter, were subject to de-multiplexing and trimming of Illumina adaptor residuals using Illumina’s bcl2fastq software version v2.20.0.422. RNA-seq reads were mapped onto *T. kivui* WT genome (<u>OZ020628</u>) using Geneious Prime 2024.0.5 with the built-in Geneious mapper (default parameters). After counting (Geneious, default parameters), the DESeq2^67^ plugin for Geneious was used for PCA– and differential expression analysis (default parameters). Genes were considered differentially expressed when |log_2_(FC)| > 1 and Benjamini and Hochberg adjusted p-value < 0.05.

### Cloning

The GoldenMOCS assembly system described before was used for all cloning^28^. An overview of plasmids used in this study is given in Supplementary Table S5. The DH10B and CopyCutter (for large vectors) *E. coli* strains were transformed using classical chemically induced competence.

SPF-B017, a recipient BB3^68^ *E. coli*-*T. kivui* shuttle vector, was constructed by combining several parts by BbsI Golden Gate assembly, i.e., a BsaI-compatible AD linker, the pMU131 Gram-positive replicon from BB2_pMU131^28^ (shortened to remove a BsaI site), the thermostable kanamycin resistance marker from pMU131^49^, and a ColE1 Gram-negative origin of replication.

The Δ*pyrE* vector contained *pyrE* flanking regions (1000 bp each) so that deletion results in the removal of most of *pyrE* CDS, except for the first three and last two codons. The resulting design was ordered as a gene synthesis (Twist Bioscience, CA, USA) cloned in a high copy vector.

The PyrE_KI_ plasmid was assembled in two steps. First, two homology arms upstream and downstream *pyrE* (1000 bp) were amplified separately with fusion sites A4 and 4D^69^. The upstream homology arm also contained the *pyrE* gene, and both amplicons were cloned as BB2 donors using the pMini2.0 kit (NEB, MA, USA). The resulting plasmids were cloned into an *E. coli* BB3 AD recipient vector^70^ by Golden Gate assembly using BsaI-HFv2 (NEB).

The Ech2_KI_ vector was similarly cloned in multiple steps. First, *pyrE* downstream homology arm similar as the one described above – but with CD fusion sites^69^ – was subcloned using the pMini2.0 kit (NEB). The AB vector containing the upstream homology arm, *pyrE* CDS, and the BBa_B1001 terminator was synthesized (Twist Bioscience). The BC vector containing *T. kivui* S-layer promoter^28^, an Esp3I 23 linker cassette and the terminator from the F0F1 ATPase operon (downstream TKV_RS03140) was also synthesized. The AB, BC and CD parts were assembled in SPF-B017 with Golden Gate assembly (BsaI), yielding BB3_pyrE_Esp3I. Finally, the Ech2 operon including the upstream TKV_RS12155 hypothetical CDS was amplified by PCR and cloned into BB3_pyrE_Esp3I by Golden Gate assembly using Esp3I (NEB), yielding the final ∼15 kb Ech2_KI_ vector.

### *T. kivui* transformation and genome editing

*T. kivui* was transformed as previously described^28^, with the exception that the selection step was performed in liquid culture before isolation on plates. For genome editing, linear DNA encompassing the homology regions was first obtained by PCR and residual plasmid removal (DpnI digestion). The linear template was used for transformation, selecting for 5-FOA resistance or uracil prototrophy. In the latter case, cells were directly transferred to minimal medium containing the linear DNA and incubated until growth occurred.

## Acknowledgements

The authors are indebted to Marlies Müller, Ivo van den Hurk, Dominic Uhlir, Julia Reichebner and Angeliki Sitara for excellent technical assistance, and Irma Querques for fruitful discussions.

This work was supported by the Christian Doppler Research Association and Circe Biotechnologie GmbH, Vienna, Austria. The authors acknowledge TU Wien Bibliothek for financial support through its Open Access Funding Program.

## Author contributions

Conceptualization, R.H., S.P.; Investigation, R.H., J.H., M.S.; Formal analysis, R.H., J.H., M.S., G.G.T and S.P; Software, J.H., G.G.T.; Visualization, R.H.; Funding acquisition, S.P.; Supervision, S.P.; Writing – original draft, R.H., M.S.; Writing – review & editing, R.H., J.H., M.S., G.G.T and S.P.

## Data availability

Sequencing raw data (Supplementary Table S6) have been deposited at the European Nucleotide Archive (https://www.ebi.ac.uk/ena/browser) under project accession PRJEB79489. The annotated genome sequence of CO-1 is available under accession XXXXX.

---

**Table 1:** Common mutations observed in CO-1, CO-2 and CO-3 strains.

| Position | Type | Locus tag | Effect |
| --- | --- | --- | --- |
| 625,773 | Substitution | TKV_RS07415 | D215N (GAT → AAT) |
| 1,027,204 | Substitution | Intergenic | Intergenic (T → A) |
| 1,266,481 | Substitution | TKV_RS04105 | A93P (GCT → CCT) |
| 1,963,781 | Substitution | TKV_RS10045 | D187E (GAT → GAA) |
| 1,968,524 | Substitution | TKV_RS10070 | A944D (GCT → GAT) |

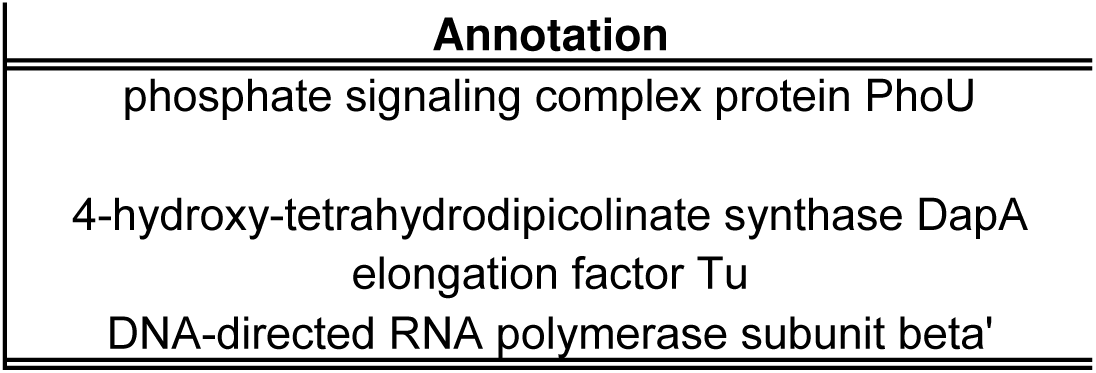

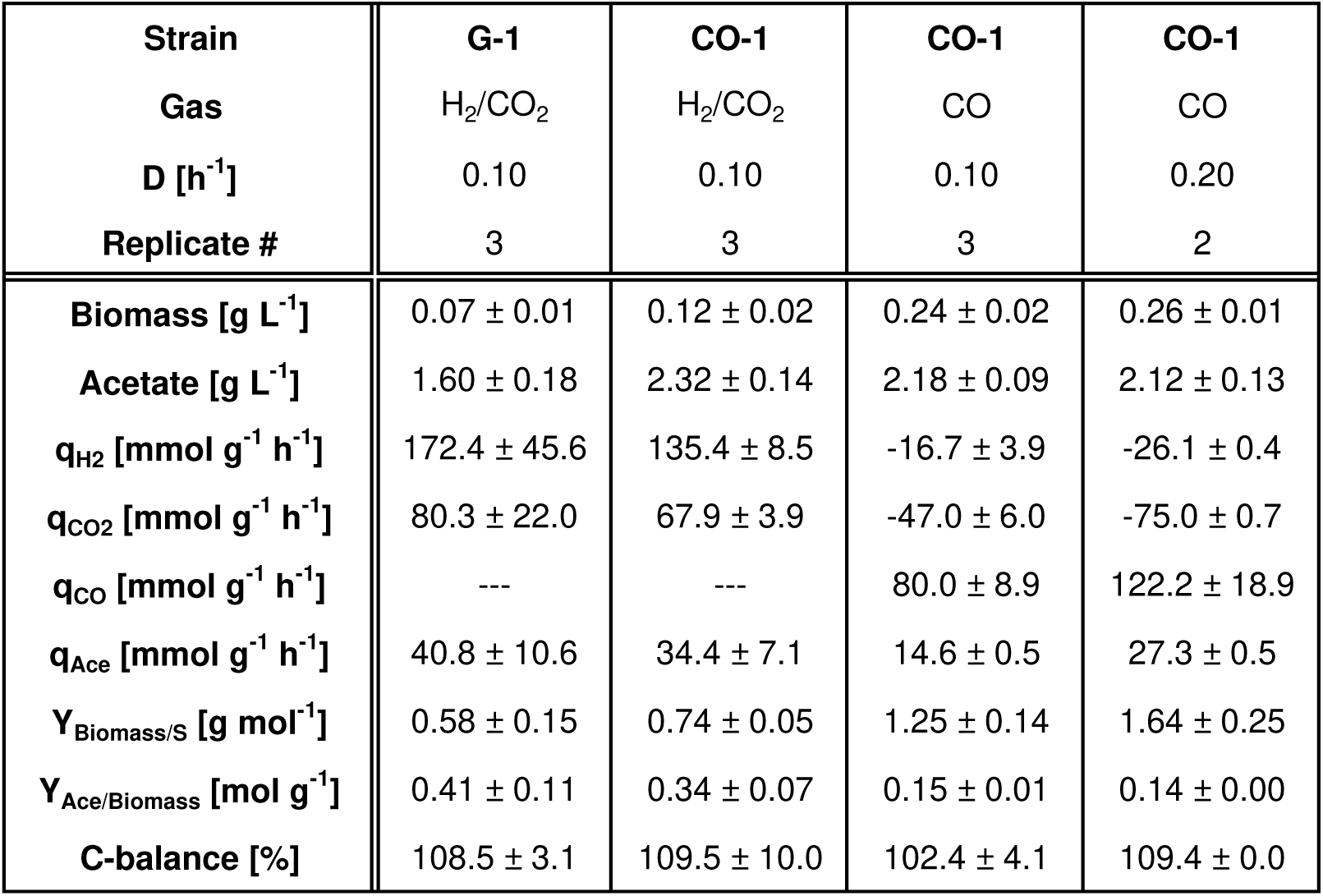

