## Supplementary figures and images for "A megatransposon drives the adaptation of *Thermoanaerobacter kivui* to carbon monoxide"

### Supplementary File 2

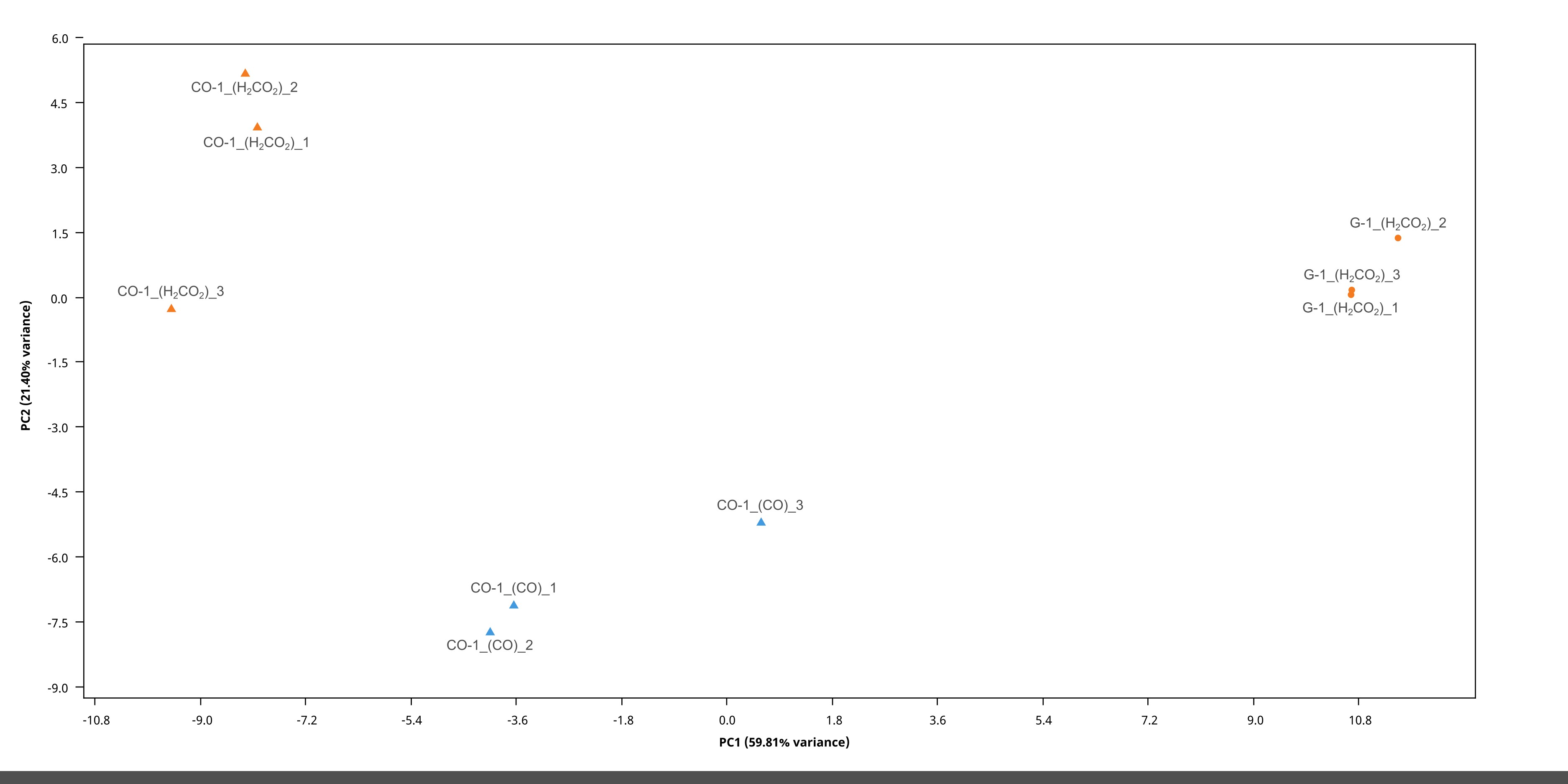
